# Simulating atmospheric drought: Silica gel packets dehumidify mesocosm microclimates

**DOI:** 10.1101/2023.10.06.561294

**Authors:** S. Varghese, B.A. Aguirre, F. Isbell, A.J. Wright

## Abstract

1. As global temperatures rise, droughts are becoming more frequent and severe. To predict how drought might affect plant communities, ecologists have traditionally designed experiments with controlled watering regimes and rainout shelters. Both treatments have proven effective for simulating soil drought. However, neither are designed to directly modify atmospheric drought.
2. Here, we detail the efficacy of a silica gel atmospheric drought treatment in outdoor mesocosms with and without a cooccurring soil drought treatment. At California State University, Los Angeles, we monitored relative humidity (RH), temperature, and vapor pressure deficit (VPD) every 10 minutes for five months in a bare-ground experiment featuring mesocosms treated with soil drought (reduced watering) and/or atmospheric drought (silica packets suspended 12 cm above soil).
3. We found that silica packets dehumidified these microclimates most effectively (-5% RH) when combined with reduced soil water, regardless of the ambient humidity levels of the surrounding air. Further, packets increased microclimate VPD most effectively (+0.4 kPa) when combined with reduced soil water and ambient air temperatures above 20°C. Finally, packets simulated atmospheric drought most consistently when replaced within three days of deployment.
4. Our results demonstrate the use of silica packets as effective dehumidification agents in outdoor drought experiments. We emphasize that incorporating atmospheric drought in existing soil drought experiments can improve our understandings of the ecological impacts of drought.

## Introduction

Climate change is driving shifts in the frequency and intensity of drought worldwide (Sheffield & Wood 2008; Trenberth 2011; Hoerling et al. 2012; Cook et al. 2014; Grossiord et al. 2020; IPCC 2021). In particular, warming is decreasing air relative humidity (RH) and increasing vapor pressure deficit (VPD), collectively driving higher evaporative demand (Burke & Brown 2008; Cook et al. 2014). While drought can be defined in many ways (Table S1; Van Loon 2015; Crausbay et al. 2017), the most widespread outcome of climate warming is *ecological drought*, or the combination of precipitation shortages and increased evaporative demand due to rising temperatures (IPCC 2021). In the United States, climate models predict that summer evaporative demand (measured using VPD) will increase by 51% by the year 2100 (Ficklin & Novick 2017).

In plant communities, drought can cause declines in species richness, increases in species extinction risk, and widespread vegetation die-off, which can have lasting impacts on ecosystem dynamics (Tilman & El Haddi 1992; Breshears et al. 2005; Allen et al. 2015; McDowell et al. 2008). Plants require water to perform basic metabolism. Importantly, relative water balance within a plant is the result of both water intake from the soil *and* water loss at the leaf surface (Schweiger et al. 2023). The later occurs via transpiration, which plants regulate by reducing their stomatal aperture (Von Caemmerer & Baker 2007; Lin et al. 2015; McAdam & Brodribb 2015). During ecological drought, plants contract stomata in response to moisture shortages in both the soil *and* the air, which can result in reduced productivity and decreased carbon fixation at an ecosystem scale (Ocheltree et al. 2014; Grossiord et al. 2020; Fu et al. 2022; Schönbeck et al. 2022).

Soil moisture and atmospheric demand do not always change in tandem (Hanks 2012; Hillel 2012; Novick et al. 2016; Fu et al. 2022). For example, while atmospheric aridity (e.g., VPD) is expected to increase worldwide as a result of warming, climate models predict that changes in precipitation (e.g., “meteorological drought,” Table S1) will be more variable (Burke & Brown 2008; Cook et al. 2014; Yuan et al. 2019). For Southern California, one theory predicts a continuation of the meteorological drought conditions that have persisted for the last 20 years (Mann & Gleick 2015), while another theory predicts *increases* in precipitation due to shifts in late-season monsoon weather patterns (Cook & Seager 2013). Moving forward, it will become critical to assess how ecosystems might be impacted by the independent and potentially interacting effects of moisture shortages both belowground (*soil drought,* a direct result of meteorological drought) and aboveground (*atmospheric drought*, a direct result of ecological drought; Novick et al. 2016; IPCC 2021; Fu et al. 2022; Table S1).

Traditionally, outdoor drought experiments have manipulated soil moisture. This is typically done by restricting soil water input as a drought treatment (at mesic sites) or by increasing soil water input in comparison to already occurring drought (at arid sites) (e.g., Baldini & Vannozzi 1999; Alster et al. 2013; Baéz et al. 2013; Copeland et al. 2016; Kreyling et al. 2016; Filazzola et al. 2018). While both practices can effectively approximate soil drought, they are not designed to directly modify atmospheric drought (Yahdjian & Sala 2002; Rana et al. 2004; Kreyling et al. 2016; Aguirre et al. 2021).

Combining soil drought treatments with experimental manipulations of air humidity may provide more comprehensive insights into ecological drought. For example, a study by Aguirre et al. (2021) found that grass community biomass was unaffected by soil drought when humidity was increased but was reduced by approximately 50% when soil drought occurred alongside ambient (naturally lower) air humidity. Another study conducted within this same experiment demonstrated that a focal species (*Poa secunda*) exhibited a higher root:shoot biomass ratio and lower leaf area only when soil drought was combined with naturally lower air humidity (Watson et al. 2023).

Aguirre et al. (2021) proposed the use of silica gel packets as a low-cost, low-tech means of capturing air moisture and thus maintaining atmospheric drought in pots simultaneously treated with soil drought. These authors reported a season-wide reduction in mesocosm RH (- 2.5%) in pots with silica packets, but this did not translate to a corresponding season-wide increase in VPD (Aguirre et al. 2021). Indeed, it is unclear how environmental conditions influence silica packet performance. For example, humid days (or times of day) might allow for stronger packet dehumidification effects (and VPD increases), while arid conditions may drive weaker effects (Aguirre et al. 2021). Furthermore, Aguirre et al. (2021) demonstrated that pots treated with increased soil moisture (independent of humidity manipulations) had higher RH and lower VPD, likely due to moisture exchange between the soil surface and the air (e.g. Zhou et al. 2019). Higher temperatures may intensify soil-to-air water transfer (often referred to as “land-atmosphere feedbacks”) because soil evaporation is temperature dependent. Indeed, warming has been shown to intensify soil drought at landscape scales due to increased soil surface evaporation (Dirmeyer et al. 2012; Hanks 2012; McHugh et al. 2015; Samaniego et al. 2018).

To help address these knowledge gaps and communicate the level of detail necessary to reproduce the silica-induced atmospheric drought treatment in other field experiments, we established an outdoor drought experiment in bare-ground mesocosms at California State University, Los Angeles (CSULA). Our setup combined independent soil drought and atmospheric drought manipulations. This experiment was briefly described in Aguirre et al. (2021). Here, we provide a full examination of the environmental and experimental conditions that influence RH and VPD modification by silica gel packets. Using this design, we explored the following questions:

Q1. How might packet efficacy (i.e., the degree of RH and VPD modification) vary in response to ambient air conditions (RH, temperature, and VPD), soil moisture, and time elapsed since packet deployment?
Q2. Does the degree of air microclimate modification depend upon rapid hourly fluctuations in ambient air moisture (RH and VPD) over the course of a day?
Q3. Are there lagged effects wherein packet efficacy might be dependent on the ambient RH and VPD conditions of the previous day?

## Materials and Methods

### Study site

This experiment was conducted at CSULA (34.0668 °N, 118.1684 °W) from January 2 through May 12, 2020, which spans southern California’s Mediterranean growing season. Between 1970 and 2019, this region received an average of 706.0 mm of precipitation annually (PRISM Climate Group) and 618.3 mm of rainfall during the typical wet season (November-April). Wet season rainfall varies widely (e.g., from 115.9 mm in 1977 to 1664.8 mm in 1973; PRISM Climate Group). During our 2020 growing season, precipitation at CSULA totaled 377.7 mm and temperature averaged 9.4 °C (PRISM Climate Group).

### Study design

Our setup featured nine polypropylene planter pots (40-cm height, 45-cm diameter, 57-L volume, CN-NCL, Greenhouse Megastore) filled with 47 L of soil obtained nearby from the Stunt Ranch Santa Monica Mountains Reserve in Calabasas, California (34.0939 °N, 118.6567 °W). Soil at this site has a pH of 7.0 and is composed of 47% sand, 31% silt, and 22% clay (Whelan et al. 2016). To improve drainage, soil was mixed 1:1 with quartzite sand, which was sterilized to avoid the introduction of external microbiota (Aguirre et al. 2021). Around each pot, we mounted open-top chambers (35-cm height × 45-cm diameter) using a PVC frame and 6-μm-thick greenhouse film with 92% light transmission (Fig. 1; 108658, Sun Master Pull and Cut Greenhouse Film, Growers Supply, Dyersville, Iowa, USA). This setup allowed for environmental contact from above (via solar irradiance, temperature, RH, and wind flow) while simultaneously providing an enclosure to capture the unique air microclimate within each pot.

**Figure 1.**
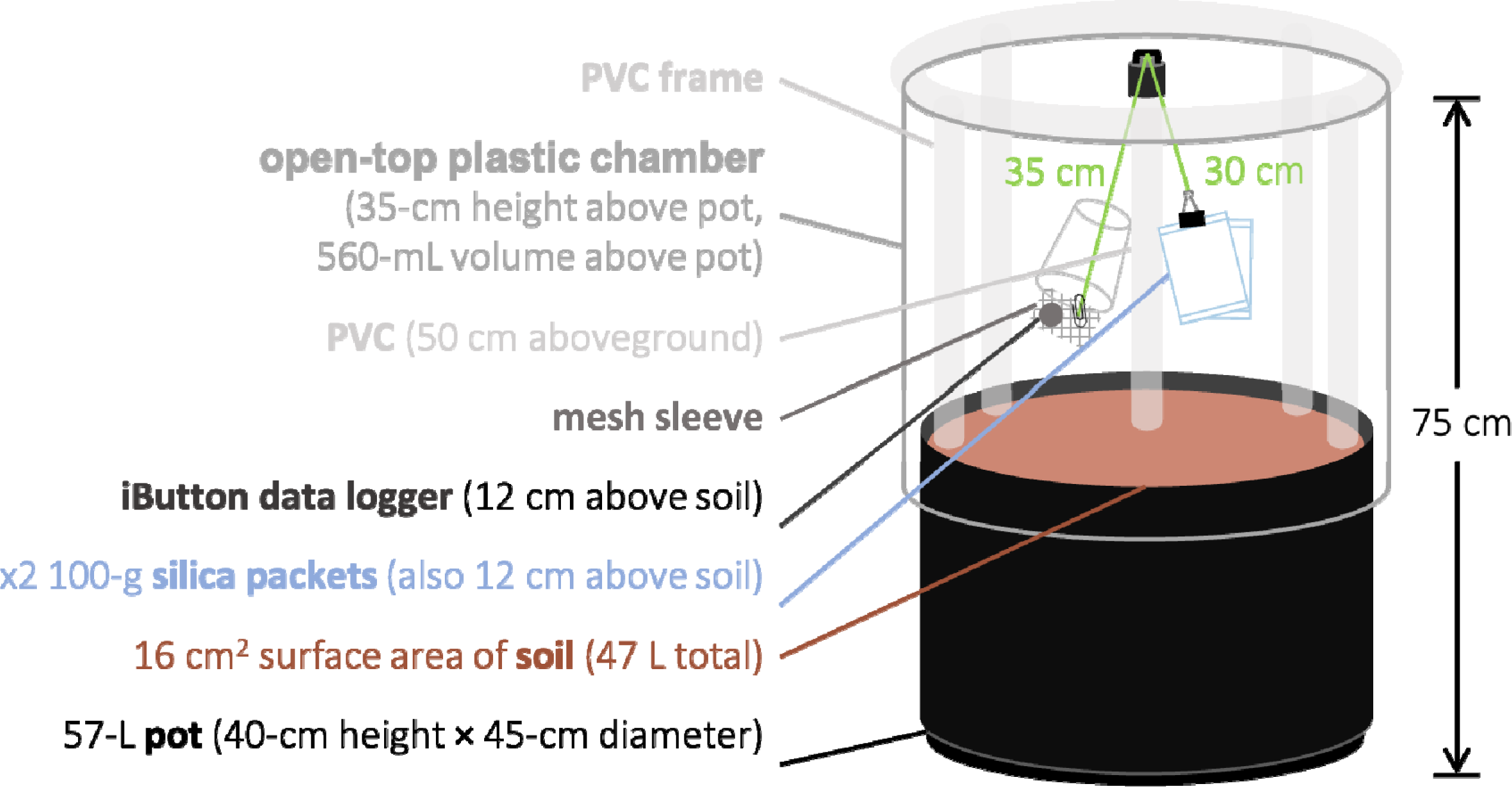
Our experiment featured nine 57-L pots, each mounted with a 35-cm-tall open-top chamber. We implemented our atmospheric drought treatment by suspending silica gel packets 12 cm above the soil surface and microclimate sensors at the same height.

Six of the nine total pots represented three true replicates of two experimental drought treatments: (1) atmospheric drought × soil drought and (2) atmospheric drought × ambient soil moisture. To impose atmospheric drought in each of our six experimental pots, we suspended two Dry & Dry 100-gram fabric silica gel packets (Dry & Dry, Brea, California, USA) 12 cm above the soil by a 30-cm string tied to a 1-m-tall PVC tube installed in the center of each pot (Fig. 1). We replaced water-saturated packets with refreshed packets every 0-7 days (see Supporting Information for our detailed protocol for packet desaturation and reuse). While we did not initially intend to vary the number of days between packet replacement, the COVID-19 pandemic had a major effect on our ability to access our experiment and facilitated an opportunity to study this variable. As a result, by the end of the study, we had high replication of replacing packets daily (54 instances × 6 pots), every other day (22 instances × 6 pots), and every two days (19 instances × 6 pots). We had less replication (≤ 8 instances) for longer durations of time between replacements.

We maintained soil moisture at either of two watering levels (ambient or drought), which we calculated based on precipitation trends at our site over the past 50 years (PRISM Climate Group). The ambient watering treatment was designed to approximate the overall average of growing season rainfall distributed evenly throughout the season (1630 mL every 4 days; Aguirre et al. 2021). The drought watering treatment fell two standard deviations below that mean (170 mL every 9 days, approximately equal to 4.8 mm annually; Aguirre et al. 2021). Upon observing unexpectedly high mortality of grasses in adjacent pots (which comprised a larger experiment), we adjusted our initial watering regime: all pots received ambient watering for 2 weeks (February 10-21), and beginning February 24, drought pots were watered 170 mL every 4 days (still 10% of the ambient volume) for the remainder of the study (Aguirre et al. 2021). Finally, during one rainy week in March, we installed a large plastic tarp to function as a rainout shelter over the entire experiment (108658, Sun Master Pull and Cut Greenhouse Film, Growers Supply, Dyersville, IA).

We maintained the remaining pots (n = 3) nearby with ambient air RH × ambient soil moisture and an identical pot and chamber setup as the six experimental pots. We averaged ambient air temperature, RH, and VPD data from these pots (which exhibited nonsignificant between-pot variability in air microclimate readings) as control reference conditions. We chose to use mesocosm-scale readings as our control reference conditions as opposed to weather station data because previous studies have shown that small-scale sensors in control pots located beside experimental pots may serve as a better reference for small-scale effects (Maclean et al. 2021). However, there are also known issues with using this type of data to make inferences about daytime air conditions (Maclean et al. 2021). Hence, we report comparisons between our mesocosm-level readings and local weather station data in Supplemental Figure S1.

### Air microclimate monitoring

In all nine pots, we installed one iButton Hygrochron datalogger (DS1923, Maxim Integrated, San Jose, CA) to monitor the air microclimate every 10 minutes. We enclosed each iButton in a mesh envelope positioned under (but not inside) a white plastic cup that blocked direct irradiance and precipitation (Wright et al. 2015). In each pot, the iButton + cup unit was tied to the central PVC pipe, allowing for the dataloggers to be suspended 12 cm above the soil surface (i.e., at the same height as the silica packets; Fig. 1). The iButtons recorded the modified air microclimate (Temp_exp_ and RH_exp_) in the experimental pots and the ambient air microclimate (Temp_amb_ and RH_amb_) in the control reference pots. We used these values to calculate pot VPD, as well (VPD_exp_ and VPD_amb_).

### Statistical modeling and analyses

Due to a temporarily malfunctioning datalogger, we excluded one pot (treated with dehumidified air × ambient soil moisture) from our analyses between March 31 and May 12. Data from the other five pots during that window, plus all six pots over the remaining dates, were averaged over hourly, 12-hour, and 24-hour intervals.

To represent the difference in RH between a pot with silica packets and a pot of ambient air, we calculated a relative humidity effect (RH_effect_) response variable as

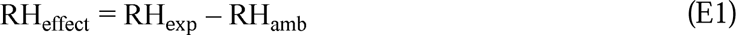

where RH_effect_ < 0 indicates microclimate dehumidification, RH_effect_ = 0 indicates no effect of packets on microclimate RH, and RH_effect_ > 0 indicates microclimate humidification. We calculated a vapor pressure deficit effect (VPD_effect_) response variable in the same way:

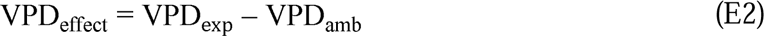

where VPD_effect_ > 0 indicates that packets dehumidified and warmed the microclimate, VPDeffect = 0 indicates no effect of packets on microclimate VPD, and VPD_effect_ < 0 indicates that packets humidified and cooled the microclimate.

Using R Statistical Software, we built linear mixed-effects models using the lmer function from the lmerTest library (Kuznetsova 2017). To investigate Q1 (How might the degree of microclimate modification vary with ambient air conditions, soil moisture, and time since packet deployment?), we focused our main analyses on the 12-hour-averaged (6:00 - 18:00) “daytime” effects as these will be most relevant for commonly studied plant behaviors (e.g., photosynthesis during daylight hours). We assigned the daytime RH_effect_ or VPD_effect_ as a continuous response variable and RH_amb_, Temp_amb_, VPD_amb_, watering treatment, count of days since packet replacement, and all higher-order interactions as fixed effects. We report on the best fit model as described below.

For Q2 (Does the degree of air microclimate modification by packets depend upon rapid hourly fluctuations in ambient air conditions over the course of a day?), we averaged the data by hour. As above, we included the daytime RH_effect_ or VPD_effect_ as a continuous response variable. We included the fixed effects of RH_amb_, Temp_amb_, VPD_amb_, and hour, as well as interactions between hour and any of the three ambient air microclimate predictors. We excluded data recorded beyond two days of packet deployment to focus on conditions for which we had higher replication. We report on the best fit model as described below.

Per Q3 (Are there lagged effects wherein packet efficacy might depend on the previous day’s air microclimate conditions?), we calculated 24-hour averages of microclimate readings and focused on cumulative two-day effects. We expect spill-over effects between days to be most straightforwardly detectable in this format. As above, we assigned the daytime RH_effect_ or VPD_effect_ as a continuous response variable. For predictors, we included the main effect of the previous day’s ambient air conditions (RH_amb_t-1__, Temp_amb_t-1__ or VPD_amb_t-1__) and its two-way interaction with the ambient air conditions of the current day, *t* (RH_amb_, Temp_amb_, or VPD_amb_). As above, we excluded all data recorded beyond two days of packet deployment. We report on the best fit model as described below.

For all models, we included date as a non-interactive fixed effect. This variable was included only to detrend our data and allow us to focus instead on the environmental and experimental influences on packet efficiency. Further, we included pot as a random effect in all models to account for repeated measurements taken in the same pots over time. Finally, we tested three different temporal autocorrelation structures (compound symmetry, first-order autoregressive, and unstructured) and consistently found that the unstructured format aligned best with our experimental design where the degrees of freedom were closest to the product of pots and dates (Crawley 2012; Isbell et al. 2015). For example, our main daytime dataset features 571 total rows of data (= (5 pots × 101 dates) + (1 pot × 66 dates)) and our unstructured daytime models both had 548 denominator degrees of freedom.

We performed a model selection analysis on each dataset (daytime, hourly, and 24-hour) and response variable (RH_effect_ and VPD_effect_). We designated the best-fit models as those with the lowest AIC. We also used df, BIC, and R^2^ to either support these decisions or identify relationships needing further testing. Finally, we analyzed the best fit model from each dataset (daytime, hourly, and 24-hour) using the anova function from the stats library (v4.2.2; R Core Team 2022).

## Results

### Q1: Daytime effects of ambient air conditions, soil watering treatments, and packet replacement

We found that the best fit model for predicting daytime dehumidification by packets included ambient air VPD, soil watering regime, days since packet replacement, and all higher order interactions (Table S2). Packet dehumidification was strongest when VPD was low (corresponding with cool, humid conditions; Table 1; Fig 2a, c, e; VPD_amb_ significant main effect; F_1, 548_ = 13.3, P = 0.0003). Overall, packets dehumidified pot microclimates within four days of deployment Table 1; Fig 3a; Days Since Replacement significant main effect; F_1, 548_ = 31.0, P < 0.0001). We also report evidence that packets dehumidified over longer periods on hot, dry days (Table 1; Fig. S2; VPD_amb_ × Days Since Replacement significant interaction; F_1, 548_ = 7.46, P = 0.01).

**Figure 2.**
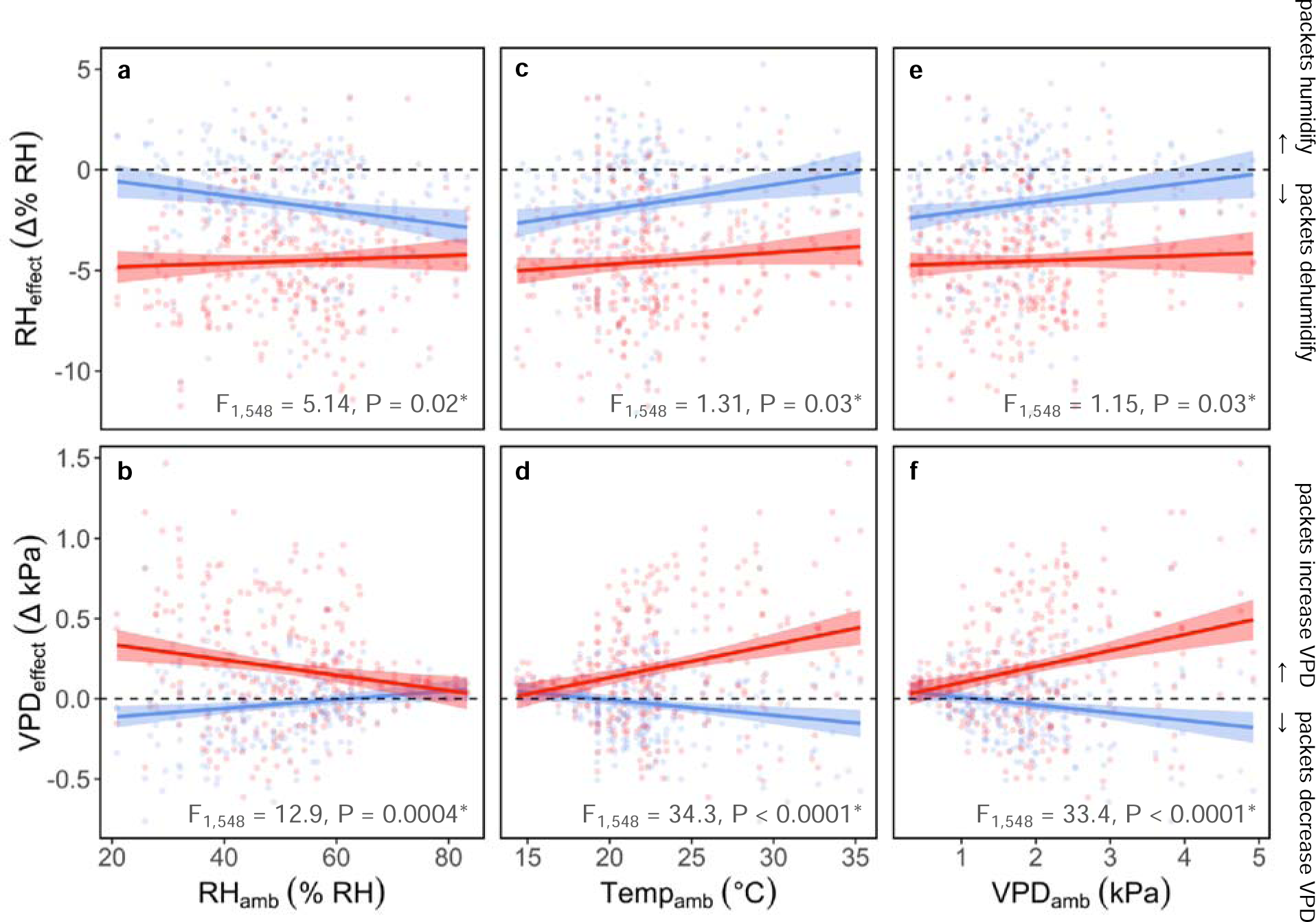
We measured changes in microclimate RH (RH_effect_) and VPD (VPD_effect_) driven by the silica gel packets in comparison with ambient pots nearby. Because VPD is a composite measure of air RH and temperature, we report on the separate roles of ambient RH (**a**-**b**) and ambient temperature (**c**-**d**), in addition to ambient VPD (**e**-**f**). Overall, packets dehumidified and increased VPD more effectively in pots of dry soil (red) than wet soil (blue). Beyond this, we found that dehumidification effects in wet-soil pots were stronger on humid days (**a**), on cool days (**c**), and when VPD was therefore lower (**e**). VPD effects were near zero in wet-soil pots (**b**, **d**, **f**). But in dry-soil pots, VPD increases were strongest on dry days (**b**), on hot days (**d**) and when VPD was therefore higher (**f**). Points represent individual dehumidified pots on separate dates, trendlines indicate significant interactions, and bands denote 95% confidence intervals.

**Figure 3.**
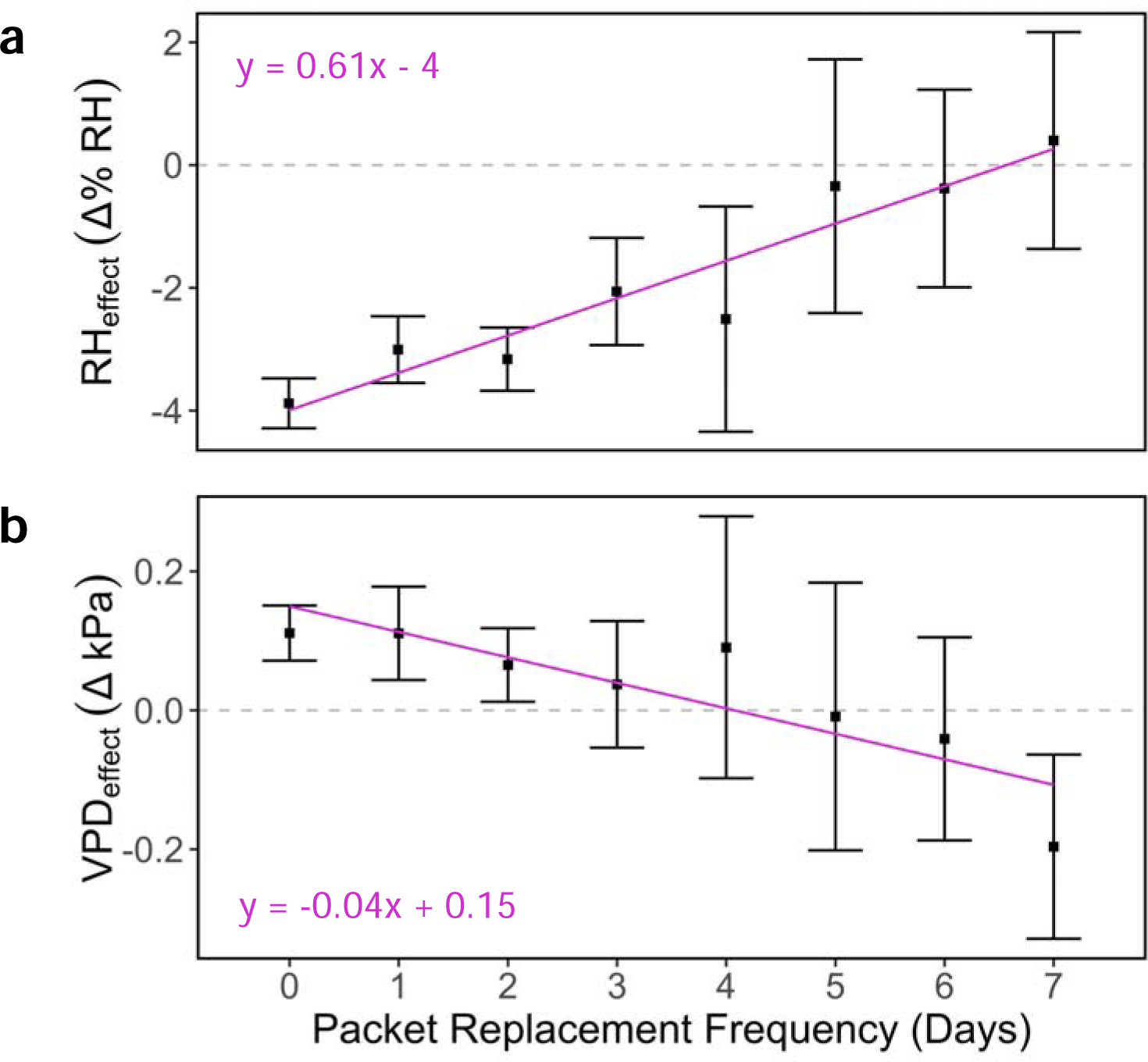
We monitored changes in RH and VPD following packet replacement (on day zero). Packets dehumidified pot microclimates within four days of deployment, beyond which dehumidification effects overlapped with zero (**a**). Packets increased microclimate VPD within two days of deployment, beyond which packet VPD effects overlapped with zero (**b**). Trendlines indicate significant relationships and error bars represent 95% confidence intervals around mean points.

**Table 1.** Our best-fit daytime models predicted the RH_effect_ and VPD_effect_ based on VPD_amb_, watering treatment, days since packet replacement, and all higher order interactions. ANOVA results significant at α = 0.05 are bolded and asterisked.

| Predictor | Fixed effects | df | F | p |
| --- | --- | --- | --- | --- |
| $RH_{\text{effect}}$ | $VPD_{\text{amb}}$ | 1, 548 | 13.3 | **0.0003\*** |
|  | Watering | 1, 6 | 4.73 | 0.07 |
|  | Days Since Replacement | 1, 548 | 31.0 | **<0.0001\*** |
|  | Date | 1, 549 | 3.08 | 0.08 |
| | $VPD_{\text{amb}} \times \text{Watering Treatment}$ | 1, 548 | 1.15 | 0.28 |
| | $VPD_{\text{amb}} \times \text{Days Since Replacement}$ | 1, 548 | 7.46 | **0.01\*** |
| | $\text{Watering Treatment} \times \text{Days Since Replacement}$ | 1, 548 | 2.19 | 0.14 |
| | $VPD_{\text{amb}} \times \text{Watering Treatment} \times \text{Days Since Replacement}$ | 1, 548 | 0.50 | 0.48 |
| $VPD_{\text{effect}}$ | $VPD_{\text{amb}}$ | 1, 548 | 0.33 | 0.57 |
|  | Watering | 1, 5 | 0.17 | 0.70 |
|  | Days Since Replacement | 1, 548 | 6.36 | **0.01\*** |
|  | Date | 1, 548 | 1.22 | 0.27 |
| | $VPD_{\text{amb}} \times \text{Watering Treatment}$ | 1, 548 | 33.4 | **<0.0001\*** |
| | $VPD_{\text{amb}} \times \text{Days Since Replacement}$ | 1, 548 | 0.41 | 0.52 |
| | $\text{Watering Treatment} \times \text{Days Since Replacement}$ | 1, 548 | 0.01 | 0.92 |
| | $VPD_{\text{amb}} \times \text{Watering Treatment} \times \text{Days Since Replacement}$ | 1, 548 | 0.14 | 0.70 |

In our corresponding analysis of daytime VPD modification by packets, we found that the same set of fixed effects (ambient VPD, watering regime, days since packet replacement, and all higher order interactions) produced one of the best-fit models within five AIC points; we report on this model here in order to make clear comparisons with the RH model (Table S2). Notably, ambient VPD interacted with our watering treatments to jointly influence packet modification of VPD: on cool, humid days, the VPD_effect_ in all pots overlapped with zero, while on hot, dry days, packets increased VPD in drought-watered pots but had a near-zero effect in ambient-watered pots (Table 1; Figure 2b, d, f; VPD_amb_ × Watering Treatment significant interaction; F_1, 548_ = 33.4 P < 0.0001). Further, packets increased pot VPD consistently within the first two days of deployment (Table 1; Fig. 3b; Days Since Replacement significant main effect; F_1, 548_ = 6.36, P = 0.01).

### Q2: Hourly packet effects

The best-fit model for predicting hourly changes in packet dehumidification included the main effects of ambient air temperature, hour of day, and their interaction term (Table S3). Packets dehumidified pot microclimates during most hours of the day, with the strength of dehumidification shifting continuously in response to diurnal fluctuations in ambient air temperature (Table 2; Fig. 4; Temp_amb_ × Hour significant interaction; F_1, 11450_ = 44.7, P < 0.0001). Dehumidification declined in the morning hours between 8:00 and 10:00 am (Fig. 4).

**Figure 4.**
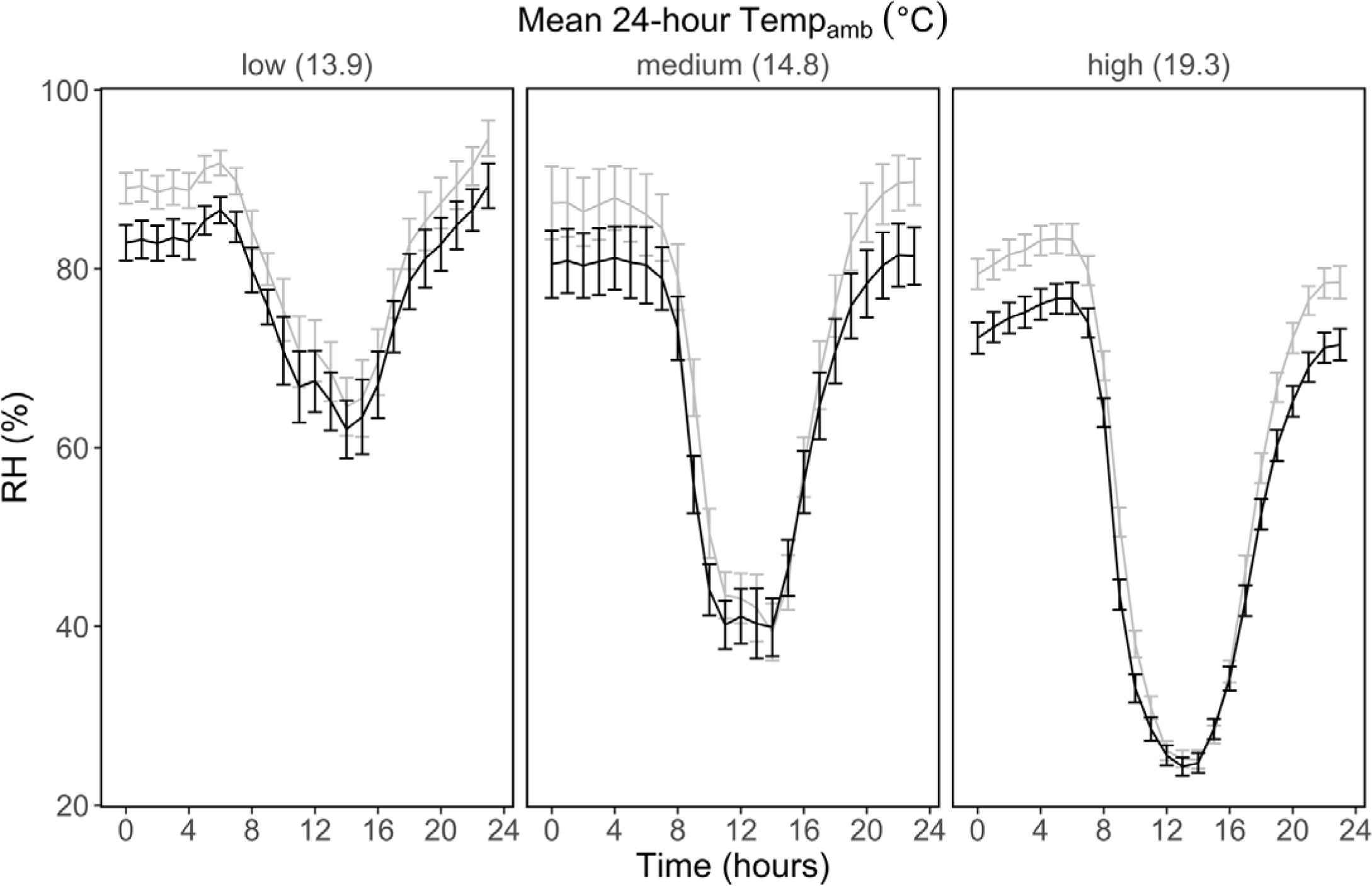
We measured fine-scale changes in RH over the course of a day. Hourly RH in pots with silica packets (black lines) can be compared with ambient air RH in nearby non-treated pots (gray lines). Both varied with respect to ambient air temperature (panels). Error bars represent 95% confidence intervals.

**Table 2.** Our best-fit hourly model predicted the RH_effect_ from the main effects and interaction of Temp_amb_ and hour. ANOVA results significant at α = 0.05 are bolded and asterisked.

| Predictor | Fixed effects | df | F | p |
| --- | --- | --- | --- | --- |
| $RH_{\text{effect}}$ | $Temp_{\text{amb}}$ | 1, 11450 | 50.2 | **<0.0001\*** |
|  | Hour | 1, 11450 | 40.1 | **<0.0001\*** |
|  | Date | 1, 11452 | 4.07 | **0.04\*** |
| | $Temp_{\text{amb}} \times \text{Hour}$ | 1, 11450 | 44.7 | **<0.0001\*** |

The best-fit model for predicting hourly changes in packet VPD modification included the main effects of ambient VPD, hour of day, and their interaction term, though because multiple models had similar AIC scores and all models had low R^2^ values (<0.09), we do not report further on those results here (Table S3; Table S4; Figure S3).

### Q3: Two-day lagged packet effects

We found that the best-fit model for predicting a two-day lagged RH_effect_ included VPD_amb_ and VPD_amb_t-1__ (Table S5) whose interaction was nonsignificant (Table S6; VPD_amb_t-1__ × VPD_amb_ nonsignificant interaction; F_1, 206_ = 1.70, P = 0.19). The best-fit model for a corresponding two-day lagged VPD_effect_ included the same predictors (VPD_amb_ and VPD_amb_t-1__, Table S5) whose interaction was again nonsignificant (Table S6; VPD_amb_t-1__ × VPD_amb_ nonsignificant interaction; F_1, 206_ = 2.48, P = 0.12).

## Discussion

In our outdoor mesocosm experiment, we found that silica packets dried air microclimates by decreasing RH and increasing VPD most effectively when soil moisture was low. The 5% RH reduction capacity we observed was sufficient to increase VPD by up to 0.4 kPa (a value similar to 50-year projections for VPD, Ficklin & Novick 2017). We also found that packets dehumidified most reliably when replaced prior to their saturation with captured moisture (see *Supporting Information* for our protocol for desaturating used packets for redeployment). As such, replacing packets within two days helped to maintain drought-level VPD at our Los Angeles field site. We found contrasting effects of ambient daily VPD on microclimate RH and VPD, indicating areas for future experimentation that we explore below. We also demonstrate that there are tradeoffs in this approach: packets likely cannot be deployed at very large scales or at remote sites that are challenging to access regularly. As such, future work should examine high-precision, automated feedback systems for maintaining outdoor VPD and RH at optimum ranges.

### Packet performance relative to soil moisture and air conditions

In our arid Mediterranean climate, we found that packets reduced ambient air RH by 2-3% in pots of wet soil when ambient VPD was lower than 2.5 kPa. Because reporting on limits for VPD is not immediately intuitive for applying these results broadly, we thus report on temperature and humidity optima, as well: packets dehumidified pots of wet soil when pot RH was above 45% and ambient pot temperatures were below 25 °C. These microclimate readings corresponded with weather station readings of 33.5% RH and 33.5 °C (Fig. S1). But when packets were combined with soil drought, they reduced pot RH by 5% regardless of ambient air temperature, relative humidity, or VPD. Similarly, packets increased pot VPD more reliably in drought-watered pots than ambient-watered pots; this VPD_effect_ was strongest (up to 0.4 kPa) on days of high VPD (and thus high atmospheric temperatures and low humidity). This evidence suggests that packet efficacy is related to stocks and flows of water in these mesocosms (e.g. Fig. S4). While packets can be used to remove water from the air, this air moisture might be rapidly replaced by evaporated soil surface water if soil moisture is high (Fig. S4b). But in drier soils, soil moisture may not be high enough to drive evaporation into dry air, which would allow us to detect stronger reductions in atmospheric humidity in dry pots (Fig. S4a).

Importantly, we also present evidence of an apparent opposition between how silica packets modify microclimate RH versus VPD under the context of shifting ambient VPD: daytime increases in ambient VPD corresponded with a stronger VPD_effect_ but a weaker RH_effect_. We speculate this trend to be the result of packets’ different physical responses to changes in humidity and temperature. In particular, packets may dehumidify more effectively with increasing ambient humidity due to passive concentration gradients: higher humidity may cause greater diffusion of water molecules towards the silica desiccant. At the same time, packets may capture moisture more effectively at higher temperatures when rates of water transfer increase. Warming drives higher rates of evaporation and soil absorption (Walker 1994; Dai et al. 2004; Smith et al. 2008; Huntington 2006; Dirmeyer et al. 2012; Hanks 2012; Samaniego et al. 2018). We thus present some clues that packets may dehumidify best when ambient air is hot and humid, though this was impossible to test directly in our arid Mediterranean climate where all of the hottest days were also the driest days. To address this gap, we recommend that future studies apply desiccation packets in tropical systems.

The VPD modification we report here (+0.4 kPa) corresponds with meaningful ecological differences. For example, Ficklin and Novick (2017) report on VPD differences of approximately 0.5 kPa between central Minnesota and arid regions of Kansas. Past work has emphasized the role that RH and VPD changes at these magnitudes can play for plant performance. For example, Grossiord et al. (2020) report that VPD changes ≥ 0.1 kPa can decrease steady-state stomatal aperture, stomatal conductance, and CO_2_ assimilation rate. Another study by Schönbeck et al. (2022) found that VPD levels as low as 1.4 kPa can lead to losses of stem conductivity, leaf water potential, and biomass, even when soil water is not limiting.

### Packets as rapid feedback systems

We found that packets can be sensitive to both immediate and cumulative changes in air conditions. For example, in our data, packets dehumidified pot microclimates from 6:00 pm to 6:00 am (overnight, when humidity was most abundant), emitted humidity around 8:00 am (possibly from oversaturation with nighttime humidity), and then returned to dehumidification at 9:00 am. This switch may have been driven by increases in water transfer rates at sunrise when the air warmed. Simultaneously, daytime warming likely expedited the evaporation of soil surface moisture, creating increased humidity in pots of wet soils especially. As such, mornings at our site are correlated with reduced efficiency of RH effects but higher efficiency of VPD effects.

Past plant physiological studies have shown that stomatal pore aperture and gas exchange operate at very fine-scale time intervals (McAdam & Brodribb 2015; Grossiord 2020). For example, Lawson & Blatt (2014) demonstrated evidence of stomata responding to sun flecks in as little as ten seconds. Understanding fast-acting physiological responses like these under the context of drought requires the implementation of a rapid-response drought manipulation. Our results suggest that silica packets can perform this level of rapid modification.

### Packet dehumidification on cumulative days following deployment

In our assessment of cumulative two-day packet effects, we found no evidence that the previous day’s ambient air VPD impacted packet VPD modification on the present day. This aligns with our finding that packets can reduce air humidity to atmospheric drought levels within the first three days of deployment. Taken together, both results suggest that packets can be deployed for consecutive days and still retain their capacity to reduce atmospheric humidity and increase atmospheric VPD.

### Study Limitations

Despite low replication of experimental units (3 pots per soil moisture level), we found robust and consistent effects of the desiccation packets. Our ability to detect these effects, despite a small number of pots, may partly be due to the high temporal resolution of humidity and temperature data collected in our pots (measurements every 10 minutes), which is higher resolution than in many other drought experiments (e.g., Adair et al. 2011; Vogel et al. 2013; Leimer et al. 2014; Wright et al. 2015; Wang et al. 2016; Fischer et al. 2019). Due to budget and time constraints, there is often a tradeoff between measuring soil moisture either more frequently across a smaller number of experimental units (as in our study) or less frequently across a larger number of experimental units (as in many other drought studies). Here, we find overwhelmingly consistent effects over time. We encourage future studies to implement higher replication, which could increase the precision of these estimated treatment effects. But we do not expect that increasing replication would change the direction or significance of these effects.

While this experiment was performed in bare-ground mesocosms, future studies that seek to implement an atmospheric drought treatment using silica gel packets, especially when vegetation is present, should consider the challenges of scaling up this approach to field experiments. For example, in the context of large, open-field plots (in comparison to mesocosms), there may be greater mixing of air across plot edges. Specifically, if the air on one side of a plot edge becomes substantially more humid (due to treatment effects or natural environmental fluctuations), the humidity may diffuse and equilibrate beyond the plot margin.

Better understanding the full mesocosm water cycle will also help apply these results more broadly (e.g. Figure S4). Further, the results presented here may differ when vegetation is present as plants connect soil moisture to air humidity through feedbacks in both transpiration and shading, which can affect microclimate temperature and rates of evaporation (Wright et al. 2014). Finally, tracking how silica dehumidification (or industrial forms of dehumidification) may drive increased rates of soil moisture loss will be an important next step in the design of atmospheric drying experiments.

### Simulating future atmospheric drying

Atmospheric drought will become more common and more severe as climate change continues to occur. Overlooking the impacts of atmospheric drying during drought manipulations can underestimate the biological mechanisms that drive drought resistance in plant communities (Ocheltree et al. 2014; Grossiord et al. 2020; Aguirre et al. 2021). To advance our understandings of how drought events will impact plant communities, we must conduct drought experiments with both soil and air drying regimes that accurately simulate natural ecological drought, which is characterized by moisture shortages at both the soil and atmospheric levels (Novick et al. 2016; Aguirre et al. 2021; IPCC 2021). This study examined the efficacy of silica gel packets as one such dehumidification solution.

## Acknowledgments

We thank all current members and alumni of the Wright Lab for their hard work in establishing and maintaining the experiment and for their assistance in collecting data. We thank John Harris for his continued assistance as our greenhouse manager. This work was supported by the La Kretz Environmental Science Endowment Grant awarded to Dr. Alexandra Wright, a Cal State Los Angeles School of Natural and Social Sciences Startup Grant awarded to Dr. Alexandra Wright, and a California State University Program for Education & Research in Biotechnology Young Investigator Grant awarded to Dr. Alexandra Wright. This work was also supported by an NSF DEB CAREER grant (#2143186). Research reported in this publication was supported by the National Institute of General Medical Sciences of the National Institutes of Health under Awards R25GM061331 and T34GM145503. The content is solely the responsibility of the authors and does not necessarily represent the official views of the National Institutes of Health. The funding for the Cal State LA LSAMP-BD Cohort XIV program is provided by the National Science Foundation under Grant HRD-1700556.

## Author Contributions

A.J.W., B.A.A., and S.V. conceived the ideas and designed the methodology; A.J.W., B.A.A., and S.V. collected data; S.V. analyzed data and A.J.W. advised on the data analysis; S.V., A.J.W., F.I., and B.A.A. contributed to interpretation of results; S.V. and A.J.W. led the writing of the manuscript. All authors contributed critically to the drafts and gave final approval for publication.

## Conflict of Interest Statement

The authors declare no conflicts of interest.

## Data Availability Statement

Data are organized into five separate spreadsheets publicly available at https://figshare.com/projects/Simulating_atmospheric_drought_Silica_gel_packets_effectively_d ehumidify_microclimates/137385 (Varghese et al. 2022).

## Notes

https://figshare.com/projects/Simulating_atmospheric_drought_Silica_gel_packets_effectively_dehumidify_microclimates/137385

### Competing Interest Statement

The authors have declared no competing interest.

## References

Adair, C., E., Reich, P. B., Trost, J. J., & Hobbie, S. E. (2011). Elevated CO_2_ stimulates grassland soil respiration by increasing carbon inputs rather than by enhancing soil moisture. Global Change Biology, 17(12), 3546–3563. 10.1111/j.1365-2486.2011.02484.x

Aguirre, B. A., Hsieh, B., Watson, S. J., & Wright, A. J. (2021). The experimental manipulation of atmospheric drought: Teasing out the role of microclimate in biodiversity experiments. Journal of Ecology, 109(5), 1986–1999. 10.1111/1365-2745.13595

Allen, C. D., Breshears, D. D., & McDowell, N. G. (2015). On underestimation of global vulnerability to tree mortality and forest dieLJoff from hotter drought in the Anthropocene. Ecosphere, 6(8), 1–55. 10.1890/ES15-00203.1

Alster, C. J., German, D. P., Lu, Y., & Allison, S. D. (2013). Microbial enzymatic responses to drought and to nitrogen addition in a southern California grassland. Soil Biology and Biochemistry, 64, 68–79. 10.1016/j.soilbio.2013.03.034

Báez, S., Collins, S. L., Pockman, W. T., Johnson, J. E., & Small, E. E. (2013). Effects of experimental rainfall manipulations on Chihuahuan Desert grassland and shrubland plant communities. Oecologia, 172, 1117–1127. 10.1007/s00442-012-2552-0

Baldini, Mario & Vannozzi, Gian Paolo. (1999). Yield relationships under drought in sunflower genotypes obtained from a wild population and cultivated sunflowers in rain-out shelter in large pots and field experiments. Helia, 22:81–86.

Breshears, D. D., Cobb, N. S., Rich, P. M., Price, K. P., Allen, C. D., Balice, R. G., … & Meyer, C. W. (2005). Regional vegetation die-off in response to global-change-type drought. Proceedings of the National Academy of Sciences, 102(42), 15144–15148. 10.1073/pnas.0505734102

Burke, E. J., & Brown, S. J. (2008). Evaluating uncertainties in the projection of future drought. Journal of Hydrometeorology, 9(2), 292–299. 10.1175/2007JHM929.1

Cook, B. I. & Seager, R. (2013) The response of the North American monsoon to increased greenhouse gas forcing. Journal of Geophysical Research: Atmospheres 118, 1690–1699. 10.1002/jgrd.50111

Cook, B. I., Smerdon, J. E., Seager, R., & Coats, S. (2014). Global warming and 21st century drying. Climate Dynamics, 43(9), 2607–2627. 10.1007/s00382-014-2075-y

Copeland, S. M., Harrison, S. P., Latimer, A. M., Damschen, E. I., Eskelinen, A. M., FernandezLJGoing, B., … & Thorne, J. H. (2016). Ecological effects of extreme drought on Californian herbaceous plant communities. Ecological Monographs, 86(3), 295–311. 10.1002/ecm.1218

Crawley, M. J. (2012). The R book. John Wiley & Sons.

Crausbay, S. D., Ramirez, A. R., Carter, S. L., Cross, M. S., Hall, K. R., Bathke, D. J., … & Sanford, T. (2017). Defining ecological drought for the twenty-first century. Bulletin of the American Meteorological Society, 98(12), 2543–2550. 10.1175/BAMS-D-16-0292.1

Dai, A., Trenberth, K. E., & Qian, T. (2004). A global dataset of Palmer Drought Severity Index for 1870–2002: Relationship with soil moisture and effects of surface warming. Journal of Hydrometeorology, 5(6), 1117–1130. 10.1175/JHM-386.1

Dirmeyer, P. A., Cash, B. A., Kinter III, J. L., Stan, C., Jung, T., Marx, L., … & Manganello, J. (2012). Evidence for enhanced land–atmosphere feedback in a warming climate. Journal of Hydrometeorology, 13(3), 981–995. 10.1175/JHM-D-11-0104.1

Ficklin, D. L., & Novick, K. A. (2017). Historic and projected changes in vapor pressure deficit suggest a continentalLJscale drying of the United States atmosphere. Journal of Geophysical Research: Atmospheres, 122(4), 2061–2079. 10.1002/2016JD025855

Filazzola, A., Liczner, A. R., Westphal, M., & Lortie, C. J. (2018). The effect of consumer pressure and abiotic stress on positive plant interactions are mediated by extreme climatic events. New Phytologist, 217(1), 140–150. 10.1111/nph.14778

Fischer, C., Leimer, S., Roscher, C., Ravenek, J., de Kroon, H., Kreutziger, Y., … & Hildebrandt, A. (2019). Plant species richness and functional groups have different effects on soil water content in a decadeLJlong grassland experiment. Journal of Ecology, 107(1), 127–141. 10.1111/1365-2745.13046

Fu, Z., Ciais, P., Prentice, I.C. et al. Atmospheric dryness reduces photosynthesis along a large range of soil water deficits. Nature Communications 13, 989 (2022). 10.1038/s41467-022-28652-7

Grossiord, C., Buckley, T. N., Cernusak, L. A., Novick, K. A., Poulter, B., Siegwolf, R. T., … & McDowell, N. G. (2020). Plant responses to rising vapor pressure deficit. New Phytologist, 226(6), 1550–1566. 10.1111/nph.16485

Hanks, R. J. (2012). Applied soil physics: soil water and temperature applications (Vol. 8). Springer Science & Business Media.

Hillel, D. (2012). Applications of soil physics. Elsevier.

Hoerling, M., Eischeid, J., Perlwitz, J., Quan, X., Zhang, T., & Pegion, P. (2012). On the increased frequency of Mediterranean drought. Journal of Climate, 25(6), 2146–2161. 10.1175/JCLI-D-11-00296.1

Huntington, T. G. (2006). Evidence for intensification of the global water cycle: Review and synthesis. Journal of Hydrology, 319(1-4), 83–95. 10.1016/j.jhydrol.2005.07.003

IPCC, 2021: Climate Change 2021: The Physical Science Basis. Contribution of Working Group I to the Sixth Assessment Report of the Intergovernmental Panel on Climate Change [Masson-Delmotte, V., P. Zhai, A. Pirani, S. L. Connors, C. Péan, S. Berger, N. Caud, Y. Chen, L. Goldfarb, M. I. Gomis, M. Huang, K. Leitzell, E. Lonnoy, J. B. R. Matthews, T. K. Maycock, T. Waterfield, O. Yelekçi, R. Yu and B. Zhou (eds.)]. Cambridge University Press. In Press.

Isbell, F., Craven, D., Connolly, J., Loreau, M., Schmid, B., Beierkuhnlein, C., … & Eisenhauer, N. (2015). Biodiversity increases the resistance of ecosystem productivity to climate extremes. Nature, 526(7574), 574–577. 10.1038/nature15374

Kreyling, J., Arfin Khan, M. A., Sultana, F., Babel, W., Beierkuhnlein, C., Foken, T., … & Jentsch, A. (2016). Drought effects in climate change manipulation experiments: quantifying the influence of ambient weather conditions and rain-out shelter artifacts. Ecosystems, 20(2), 301–315. 10.1007/s10021-016-0025-8

Kuznetsova A, Brockhoff PB, Christensen RHB (2017). “lmerTest Package: Tests in Linear Mixed Effects Models.” Journal of Statistical Software, 82(13), 1–26. doi:10.18637/jss.v082.i13

Lawson, T., & Blatt, M. R. (2014). Stomatal size, speed, and responsiveness impact on photosynthesis and water use efficiency. Plant Physiology, 164(4), 1556–1570. 10.1104/pp.114.237107

Leimer, S., Kreutziger, Y., Rosenkranz, S., Beßler, H., Engels, C., Hildebrandt, A., … & Wilcke, W. (2014). Plant diversity effects on the water balance of an experimental grassland. Ecohydrology, 7(5), 1378–1391. 10.1002/eco.1464

Lin, Y., Medlyn, B., Duursma, R. et al. (2015). Optimal stomatal behaviour around the world. Nature Climate Change 5, 459–464. 10.1038/nclimate2550

Maclean, I. M., & Klinges, D. H. (2021). Microclimc: A mechanistic model of above, below and within-canopy microclimate. Ecological Modelling, 451, 109567. 10.1016/j.ecolmodel.2021.109567

Mann, M. E., & Gleick, P. H. (2015). Climate change and California drought in the 21st century. Proceedings of the National Academy of Sciences, 112(13), 3858–3859. 10.1073/pnas.1503667112

McAdam, S. A., & Brodribb, T. J. (2015). The evolution of mechanisms driving the stomatal response to vapor pressure deficit. Plant Physiology, 167(3), 833–843. 10.1104/pp.114.252940

McDowell, N., Pockman, W. T., Allen, C. D., Breshears, D. D., Cobb, N., Kolb, T., … & Yepez, E. A. (2008). Mechanisms of plant survival and mortality during drought: why do some plants survive while others succumb to drought?. New phytologist, 178(4), 719–739. 10.1111/j.1469-8137.2008.02436.x

McHugh, T. A., Morrissey, E. M., Reed, S. C., Hungate, B. A., & Schwartz, E. (2015). Water from air: an overlooked source of moisture in arid and semiarid regions. Scientific Reports, 5(1), 13767. 10.1038/srep13767

Novick, K. A., Ficklin, D. L., Stoy, P. C., Williams, C. A., Bohrer, G., Oishi, A. C., … & Phillips, R. P. (2016). The increasing importance of atmospheric demand for ecosystem water and carbon fluxes. Nature Climate Change, 6(11), 1023–1027. 10.1038/nclimate3114

Ocheltree, T. W., Nippert, J. B., & Prasad, P. V. V. (2014). Stomatal responses to changes in vapor pressure deficit reflect tissueLJspecific differences in hydraulic conductance. *Plant*, Cell & Environment, 37(1), 132–139. 10.1111/pce.12137

PRISM Climate Group, Oregon State University, https://prism.oregonstate.edu, data created 8 Dec 2020, accessed 8 Dec 2020.

R Core Team (2022). R: A language and environment for statistical computing. R Foundation for Statistical Computing, Vienna, Austria. https://www.R-project.org/.

Rana, G., Katerji, N., Introna, M., & Hammami, A. (2004). Microclimate and plant water relationship of the “overhead” table grape vineyard managed with three different covering techniques. Scientia Horticulturae, 102(1), 105–120. 10.1016/j.scienta.2003.12.008

R Core Team (2022). R: A language and environment for statistical computing. R Foundation for Statistical Computing, Vienna, Austria. URL https://www.R-project.org/

RStudio Team (2021). RStudio: Integrated Development for R. RStudio, PBC, Boston, MA. http://www.rstudio.com/

Samaniego, L., Thober, S., Kumar, R., Wanders, N., Rakovec, O., Pan, M., … & Marx, A. (2018). Anthropogenic warming exacerbates European soil moisture droughts. Nature Climate Change, 8(5), 421–426. 10.1038/s41558-018-0138-5

Schönbeck, L. C., Schuler, P., Lehmann, M. M., Mas, E., Mekarni, L., Pivovaroff, A. L., … & Grossiord, C. (2022). Increasing temperature and vapour pressure deficit lead to hydraulic damages in the absence of soil drought. Plant, Cell & Environment, 45(11), 3275–3289. 10.1111/pce.14425

Schweiger, A. H., Zimmermann, T., Poll, C., Marhan, S., Leyrer, V., & Berauer, B. J. (2023). The need to decipher plant drought stress along the soil–plant–atmosphere continuum. Oikos, 2023(9), e10136. 10.1111/oik.10136

Sheffield, J., & Wood, E. F. (2008). Projected changes in drought occurrence under future global warming from multi-model, multi-scenario, IPCC AR4 simulations. Climate Dynamics, 31(1), 79-105. 10.1007/s00382-007-0340-z

Smith, P., Fang, C., Dawson, J. J., & Moncrieff, J. B. (2008). Impact of global warming on soil organic carbon. Advances in agronomy, 97, 1–43. 10.1016/S0065-2113(07)00001-6

Smith, M. N., Taylor, T. C., van Haren, J., Rosolem, R., Restrepo-Coupe, N., Adams, J., … & Saleska, S. R. (2020). Empirical evidence for resilience of tropical forest photosynthesis in a warmer world. Nature Plants, 6(10), 1225–1230. 10.1038/s41477-020-00780-2

Tilman, D., & El Haddi, A. (1992). Drought and biodiversity in grasslands. Oecologia, 89(2), 257–264. 10.1007/BF00317226

Trenberth, K. E. (2011). Changes in precipitation with climate change. Climate Research, 47(1-2), 123–138. 10.3354/cr00953

Van Loon, A. F. (2015). Hydrological drought explained. Wiley Interdisciplinary Reviews: Water, 2(4), 359–392. 10.1002/wat2.1085

Varghese, S., Aguirre, B., Isbell, F., & Wright, A. (2022): Des_24Hour.csv. Figshare. Dataset. 10.6084/m9.figshare.19611819.v1

Varghese, S., Aguirre, B., Isbell, F., & Wright, A. (2022): Des_DayNight.csv. Figshare. Dataset. 10.6084/m9.figshare.19611822.v1

Varghese, S., Aguirre, B., Isbell, F., & Wright, A. (2022): Des_Daytime.csv. Figshare. Dataset. 10.6084/m9.figshare.19611855.v1

Varghese, S., Aguirre, B., Isbell, F., & Wright, A. (2022): Des_Hourly_1.csv. Figshare. Dataset. 10.6084/m9.figshare.19611813.v1

Varghese, S., Aguirre, B., Isbell, F., & Wright, A. (2022): Des_Hourly_2.csv. Figshare. Dataset. 10.6084/m9.figshare.19611816.v1

Vogel, A., Eisenhauer, N., Weigelt, A., & SchererLJLorenzen, M. (2013). Plant diversity does not buffer drought effects on earlyLJstage litter mass loss rates and microbial properties. Global Change Biology, 19(9), 2795–2803. 10.1111/gcb.12225

Von Caemmerer, S., & Baker, N. (2007). The biology of transpiration. From guard cells to globe. Plant Physiology, 143(1), 3–3. 10.1104/pp.104.900213

Wang, R., Gamon, J. A., Montgomery, R. A., Townsend, P. A., Zygielbaum, A. I., Bitan, K., … & Cavender-Bares, J. (2016). Seasonal variation in the NDVI–species richness relationship in a prairie grassland experiment (Cedar Creek). Remote Sensing, 8(2), 128. 10.3390/rs8020128

Walker, B. H. (1994). Landscape to regional-scale responses of terrestrial ecosystems to global change. Ambio (Journal of the Human Environment, Research and Management);(Sweden), 23(1).

Watson, S. J., Aguirre, B. A., & Wright, A. J. (2023). Soil versus atmospheric drought: A test case of plant functional trait responses. Ecology, e4109. 10.1002/ecy.4109

Whelan, M. E., Hilton, T. W., Berry, J. A., Berkelhammer, M., Desai, A. R., & Campbell, J. E. (2016). Carbonyl sulfide exchange in soils for better estimates of ecosystem carbon uptake. Atmospheric Chemistry and Physics, 16(6), 3711–3726. 10.5194/acp-16-3711-2016

Wright, A., Schnitzer, S. A., & Reich, P. B. (2014). Living close to your neighbors: the importance of both competition and facilitation in plant communities. Ecology, 95(8), 2213–2223. 10.1890/13-1855.1

Wright, A., Schnitzer, S. A., & Reich, P. B. (2015). Daily environmental conditions determine the competition–facilitation balance for plant water status. Journal of Ecology, 103(3), 648–656. 10.1111/1365-2745.12397

Yahdjian, L., & Sala, O. E. (2002). A rainout shelter design for intercepting different amounts of rainfall. Oecologia, 133(2), 95–101. 10.1007/s00442-002-1024-3

Yuan, W., Zheng, Y., Piao, S., Ciais, P., Lombardozzi, D., Wang, Y., … & Yang, S. (2019). Increased atmospheric vapor pressure deficit reduces global vegetation growth. Science advances, 5(8), eaax1396. 10.1126/sciadv.aax1396

Zhou, S., Williams, A. P., Berg, A. M., Cook, B. I., Zhang, Y., Hagemann, S., … & Gentine, P. (2019). Land–atmosphere feedbacks exacerbate concurrent soil drought and atmospheric aridity. Proceedings of the National Academy of Sciences, 116(38), 18848–18853. 10.1073/pnas.1904955116

