## Appendix for "Simulating atmospheric drought: Silica gel packets dehumidify mesocosm microclimates"

**Running head**: Simulating atmospheric drought with silica gel packets

Varghese, S.^1,2^*, Aguirre, B.A.^3^, Isbell, F.^2^, Wright, A.J.^1^

^1^ California State University Los Angeles, Department of Biological Sciences, Los Angeles, CA

^2^ University of Minnesota, Department of Ecology, Evolution, and Behavior, Minneapolis, MN

^3^ Cornell University, Department of Ecology and Evolutionary Biology, Ithaca, NY

**Corresponding author**

* Steph Varghese

**Co-author emails**

Beatriz A. Aguirre

Forest Isbell

Alexandra J. Wright

**Table S1.** Defining different types of drought according to IPCC 2021 (supported in Van Loon 2015; Crausbay et al. 2017).

| **Drought Term** | **Definition** | **Real-World Impacts** |
| --- | --- | --- |
| meteorological drought | a shortage of precipitation | reduced soil moisture; hydrological drought, agricultural drought, and socioeconomic drought |
| hydrological drought | a deficit in streamflow | below-normal levels in groundwater, lakes, and reservoirs; declining wetland area; decreased river discharge |
| agricultural drought | a shortage of soil moisture that directly impacts crop production during the growing season | reduced crop quality or yield |
| ecological drought | the result of the combination of precipitation shortages and increased evaporative demand | ecosystems are driven beyond thresholds of vulnerability, resulting in cascading negative impacts on ecosystem services |

**Table S2.** Model selection results for our daytime dataset. All models included Date as a non-interactive fixed effect and a random effect of Pot. RH_effect_ describes the difference in RH between silica-dehumidified mesocosms and ambient air conditions (RH_amb_). Similarly, Temp_amb_ = ambient temperature and VPD_amb_ = ambient vapor pressure deficit. The best fit model is indicated with an asterisk.

| **Response** | **Predictors** | **df** | **AIC** | **BIC** | **conditional R^2^** |
| --- | --- | --- | --- | --- | --- |
| RH_effect_ | RH_amb_ | 5 | 2734 | 2756 | 0.439 |
|  | Temp_amb_ | 5 | 2733 | 2755 | 0.439 |
|  | VPD_amb_ | 5 | 2731 | 2753 | 0.437 |
|  | Watering Treatment | 5 | 2722 | 2744 | 0.423 |
|  | Days Since Replacement | 5 | 2636 | 2658 | 0.482 |
|  | RH_amb_ × Temp_amb_ | 7 | 2753 | 2784 | 0.438 |
|  | RH_amb_ × VPD_amb_ | 7 | 2743 | 2774 | 0.440 |
|  | RH_amb_ × Watering Treatment | 7 | 2729 | 2759 | 0.433 |
|  | RH_amb_ × Days Since Replacement | 7 | 2645 | 2675 | 0.491 |
|  | Temp_amb_ × VPD_amb_ | 7 | 2741 | 2771 | 0.444 |
|  | Temp_amb_ × Watering Treatment | 7 | 2728 | 2759 | 0.437 |
|  | Temp_amb_ × Days Since Replacement | 7 | 2631 | 2661 | 0.500 |
|  | VPD_amb_ × Watering Treatment | 7 | 2725 | 2755 | 0.430 |
|  | VPD_amb_ × Days Since Replacement | 7 | 2633 | 2663 | 0.491 |
|  | Watering Treatment × Days Since Replacement | 7 | 2630 | 2661 | 0.474 |
|  | RH_amb_ × Temp_amb_ × VPD_amb_ | 11 | 2746 | 2794 | 0.475 |
|  | RH_amb_ × Temp_amb_ × Watering Treatment | 11 | 2765 | 2813 | 0.434 |
|  | RH_amb_ × Temp_amb_ × Days Since Replacement | 11 | 2675 | 2722 | 0.499 |
|  | RH_amb_ × VPD_amb_ × Watering Treatment | 11 | 2749 | 2796 | 0.432 |
|  | RH_amb_ × VPD_amb_ × Days Since Replacement | 11 | 2666 | 2713 | 0.493 |
|  | RH_amb_ × Watering Treatment × Days Since Replacement | 11 | 2649 | 2697 | 0.489 |
|  | Temp_amb_ × VPD_amb_ × Watering Treatment | 11 | 2743 | 2791 | 0.441 |
|  | Temp_amb_ × VPD_amb_ × Days Since Replacement | 11 | 2651 | 2699 | 0.501 |
|  | Temp_amb_ × Watering Treatment × Days Since Replacement | 11 | 2629 | 2677 | 0.506 |
|  | VPD_amb_ × Watering Treatment × Days Since Replacement | 11 | 2628* | 2676 | 0.491 |
|  | RH_amb_ × Temp_amb_ × VPD_amb_ × Watering Treatment | 19 | 2767 | 2849 | 0.488 |
|  | RH_amb_ × Temp_amb_ × VPD_amb_ × Days Since Replacement | 19 | 2707 | 2789 | 0.522 |
|  | RH_amb_ × Temp_amb_ × Watering Treatment × Days Since Replacement | 19 | 2709 | 2792 | 0.504 |
|  | RH_amb_ × VPD_amb_ × Watering Treatment × Days Since Replacement | 19 | 2687 | 2769 | 0.495 |
|  | Temp_amb_ × VPD_amb_ × Watering Treatment × Days Since Replacement | 19 | 2665 | 2747 | 0.508 |
|  | RH_amb_ × Temp_amb_ × VPD_amb_ × Watering Treatment × Days Since Replacement | 35 | 2772 | 2924 | 0.540 |
| VPD_effect_ | RH_amb_ | 5 | 112 | 133 | 0.432 |
|  | Temp_amb_ | 5 | 104 | 126 | 0.435 |
|  | VPD_amb_ | 5 | 101 | 123 | 0.436 |
|  | Watering Treatment | 5 | 102 | 124 | 0.454 |
|  | Days Since Replacement | 5 | 77 | 99 | 0.457 |
|  | RH_amb_ × Temp_amb_ | 7 | 134 | 164 | 0.435 |
|  | RH_amb_ × VPD_amb_ | 7 | 124 | 154 | 0.439 |
|  | RH_amb_ × Watering Treatment | 7 | 104 | 135 | 0.476 |
|  | RH_amb_ × Days Since Replacement | 7 | 105 | 136 | 0.458 |
|  | Temp_amb_ × VPD_amb_ | 7 | 123 | 153 | 0.435 |
|  | Temp_amb_ × Watering Treatment | 7 | 59 | 90 | 0.527 |
|  | Temp_amb_ × Days Since Replacement | 7 | 95 | 125 | 0.462 |
|  | VPD_amb_ × Watering Treatment | 7 | 57 | 88 | 0.518 |
|  | VPD_amb_ × Days Since Replacement | 7 | 91 | 121 | 0.461 |
|  | Watering Treatment × Days Since Replacement | 7 | 86 | 117 | 0.482 |
|  | RH_amb_ × Temp_amb_ × VPD_amb_ | 11 | 161 | 209 | 0.452 |
|  | RH_amb_ × Temp_amb_ × Watering Treatment | 11 | 117 | 165 | 0.527 |
|  | RH_amb_ × Temp_amb_ × Days Since Replacement | 11 | 153 | 201 | 0.463 |
|  | RH_amb_ × VPD_amb_ × Watering Treatment | 11 | 102 | 150 | 0.526 |
|  | RH_amb_ × VPD_amb_ × Days Since Replacement | 11 | 140 | 188 | 0.464 |
|  | RH_amb_ × Watering Treatment × Days Since Replacement | 11 | 119 | 166 | 0.502 |
|  | Temp_amb_ × VPD_amb_ × Watering Treatment | 11 | 94 | 142 | 0.528 |
|  | Temp_amb_ × VPD_amb_ × Days Since Replacement | 11 | 129 | 177 | 0.464 |
|  | Temp_amb_ × Watering Treatment × Days Since Replacement | 11 | 62 | 109 | 0.556 |
|  | VPD_amb_ × Watering Treatment × Days Since Replacement | 11 | 58 | 105 | 0.544 |
|  | RH_amb_ × Temp_amb_ × VPD_amb_ × Watering Treatment | 19 | 170 | 252 | 0.555 |
|  | RH_amb_ × Temp_amb_ × VPD_amb_ × Days Since Replacement | 19 | 233 | 316 | 0.472 |
|  | RH_amb_ × Temp_amb_ × Watering Treatment × Days Since Replacement | 19 | 176 | 258 | 0.560 |
|  | RH_amb_ × VPD_amb_ × Watering Treatment × Days Since Replacement | 19 | 153 | 235 | 0.554 |
|  | Temp_amb_ × VPD_amb_ × Watering Treatment × Days Since Replacement | 19 | 127 | 210 | 0.561 |
|  | RH_amb_ × Temp_amb_ × VPD_amb_ × Watering Treatment × Days Since Replacement | 35 | 326 | 477 | 0.577 |

**Table S3.** Model selection results for our hourly dataset. All models included Date as a non-interactive fixed effect and a random effect of Pot. RH_effect_ describes the difference in RH between silica-dehumidified mesocosms and ambient air conditions (RH_amb_). Similarly, Temp_amb_ = ambient temperature and VPD_amb_ = ambient vapor pressure deficit. The best fit model is indicated with an asterisk.

| **Response** | **Predictors** | **df** | **AIC** | **BIC** | **conditional R^2^** |
| --- | --- | --- | --- | --- | --- |
| RH_effect_ | Hour | 5 | 70742 | 70779 | 0.178 |
|  | RH_amb_ + Hour | 6 | 69007 | 69051 | 0.292 |
|  | Temp_amb_ + Hour | 6 | 68932 | 68976 | 0.300 |
|  | VPD_amb_ + Hour | 6 | 69165 | 69209 | 0.283 |
|  | RH_amb_ × Hour | 7 | 68971 | 69023 | 0.296 |
|  | Temp_amb_ × Hour | 7 | 68900* | 68952 | 0.303 |
|  | VPD_amb_ × Hour | 7 | 69170 | 69221 | 0.284 |
| VPD_effect_ | Hour | 5 | 15856 | 15893 | 0.081 |
|  | RH_amb_ + Hour | 6 | 15869 | 15913 | 0.082 |
|  | Temp_amb_ + Hour | 6 | 15855 | 15899 | 0.083 |
|  | VPD_amb_ + Hour | 6 | 15856 | 15900 | 0.082 |
|  | RH_amb_ × Hour | 7 | 15856 | 15907 | 0.084 |
|  | Temp_amb_ × Hour | 7 | 15867 | 15918 | 0.083 |
|  | VPD_amb_ × Hour | 7 | 15865 | 15917 | 0.083 |

**Table S4.** Our best-fit hourly model predicted the VPD_effect_ from the main effects and interaction of RH_amb_ and hour. ANOVA results significant at α = 0.05 are bolded and asterisked. VPD_effect_ describes the difference in VPD between mesocosms with silica packets and ambient air conditions. RH_amb_ = ambient relative humidity.

| **Predictor** | **Fixed effects** | **df** | **F** | **p** |
| --- | --- | --- | --- | --- |
| VPD_effect_ | RH_amb_ | 1, 11450 | 21.4 | **<0.0001*** |
|  | Hour | 1, 11450 | 34.1 | **<0.0001*** |
|  | Date | 1, 11454 | 6.07 | **0.01*** |
|  | RH_amb_ × Hour | 1, 11450 | 34.3 | **<0.0001*** |

**Table S5.** Model selection results for our 24-hour dataset. All models included Date as a non-interactive fixed effect and a random effect of Pot. RH_effect_ describes the difference in RH between silica-dehumidified mesocosms and ambient air conditions (RH_amb_). Similarly, Temp_amb_ = ambient temperature and VPD_amb_ = ambient vapor pressure deficit. The best fit model involving an interaction is indicated with an asterisk.

| **Response** | **Predictors** | **df** | **AIC** | **BIC** | **conditional R^2^** |
| --- | --- | --- | --- | --- | --- |
| RH_effect_ | $\text{RH}_{\text{amb}_{\text{t-1}}}$ | 5 | 987 | 1003 | 0.434 |
|  | $\text{Temp}_{\text{amb}_{\text{t-1}}}$ | 5 | 981 | 998 | 0.436 |
|  | $\text{VPD}_{\text{amb}_{\text{t-1}}}$ | 5 | 976 | 993 | 0.445 |
|  | RH_amb_ × $\text{RH}_{\text{amb}_{\text{t-1}}}$ | 7 | 1007 | 1031 | 0.434 |
|  | RH_amb_ × $\text{Temp}_{\text{amb}_{\text{t-1}}}$ | 7 | 997 | 1020 | 0.445 |
|  | RH_amb_ × $\text{VPD}_{\text{amb}_{\text{t-1}}}$ | 7 | 990 | 1014 | 0.446 |
|  | Temp_amb_ × $\text{Temp}_{\text{amb}_{\text{t-1}}}$ | 7 | 989 | 1012 | 0.456 |
|  | Temp_amb_ × $\text{VPD}_{\text{amb}_{\text{t-1}}}$ | 7 | 984 | 1007 | 0.455 |
|  | VPD_amb_ × $\text{VPD}_{\text{amb}_{\text{t-1}}}$ | 7 | 979* | 1003 | 0.446 |
| VPD_effect_ | $\text{RH}_{\text{amb}_{\text{t-1}}}$ | 5 | -154 | -138 | 0.402 |
|  | $\text{Temp}_{\text{amb}_{\text{t-1}}}$ | 5 | -174 | -157 | 0.442 |
|  | $\text{VPD}_{\text{amb}_{\text{t-1}}}$ | 5 | -172 | -155 | 0.432 |
|  | RH_amb_ × $\text{RH}_{\text{amb}_{\text{t-1}}}$ | 7 | -128 | -104 | 0.415 |
|  | RH_amb_ × $\text{Temp}_{\text{amb}_{\text{t-1}}}$ | 7 | -147 | -123 | 0.450 |
|  | RH_amb_ × $\text{VPD}_{\text{amb}_{\text{t-1}}}$ | 7 | -148 | -124 | 0.437 |
|  | Temp_amb_ × $\text{Temp}_{\text{amb}_{\text{t-1}}}$ | 7 | -152 | -128 | 0.450 |
|  | Temp_amb_ × $\text{VPD}_{\text{amb}_{\text{t-1}}}$ | 7 | -154 | -130 | 0.439 |
|  | VPD_amb_ × $\text{VPD}_{\text{amb}_{\text{t-1}}}$ | 7 | -159* | -135 | 0.437 |

**Table S6.** Our best-fit 24-hour model predicted the RH_effect_ and VPD_effect_ from the main effects and interaction of VPD_amb_ and $\text{VPD}_{\text{amb}_{\text{t-1}}}$. ANOVA results significant at α = 0.05 are bolded and asterisked. RH_effect_ describes the difference in RH between silica-dehumidified mesocosms and ambient air conditions. VPD_amb_ = ambient vapor pressure deficit and $\text{VPD}_{\text{amb}_{\text{t-1}}}$ = ambient vapor pressure deficit on the previous day.

| **Predictor** | **Fixed effects** | **df** | **F** | **p** |
| --- | --- | --- | --- | --- |
| RH_effect_ | VPD_amb_ | 1, 206 | 1.28 | 0.26 |
|  | $\text{VPD}_{\text{amb}_{\text{t-1}}}$ | 1, 206 | 0.14 | 0.71 |
|  | Date | 1, 207 | 6.27 | **0.01*** |
|  | VPD_amb_ × $\text{VPD}_{\text{amb}_{\text{t-1}}}$ | 1, 206 | 1.70 | 0.19 |
| VPD_effect_ | VPD_amb_ | 1, 206 | 0.75 | 0.39 |
|  | $\text{VPD}_{\text{amb}_{\text{t-1}}}$ | 1, 206 | 0.23 | 0.64 |
|  | Date | 1, 207 | 1.31 | 0.25 |
|  | VPD_amb_ × $\text{VPD}_{\text{amb}_{\text{t-1}}}$ | 1, 206 | 2.48 | 0.12 |

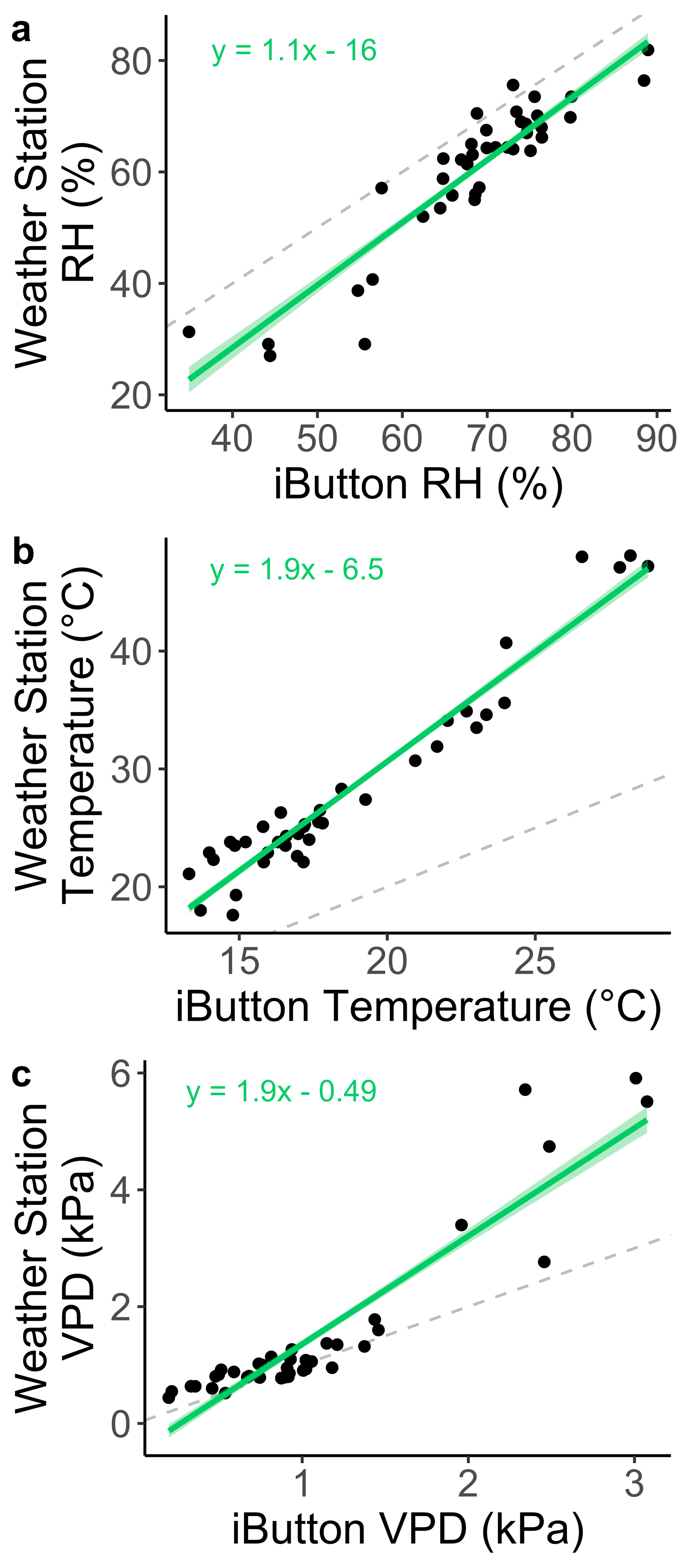

**Figure S1.** We compare pot microclimate factors (RH, temperature, and VPD) to weather station data. Although small-scale sensors in control pots located beside experimental pots may serve as a better reference for mesocosm effects, there are known issues with using this type of data to make inferences about daytime air conditions (MacLean et al. 2021). Hence, we report RH (a), temperature (b), and VPD (c) comparisons between our mesocosm-level readings (measured at CSULA located 34.0668 °N and 118.1684 °W) and local weather station data (measured at Bob Hope Airport Weather Station located 34.20045 °N and 118.35873 °W). Dashed lines represent perfect 1:1 correlation, green trendlines indicate significant relationships, and green bands denote 95% confidence intervals.

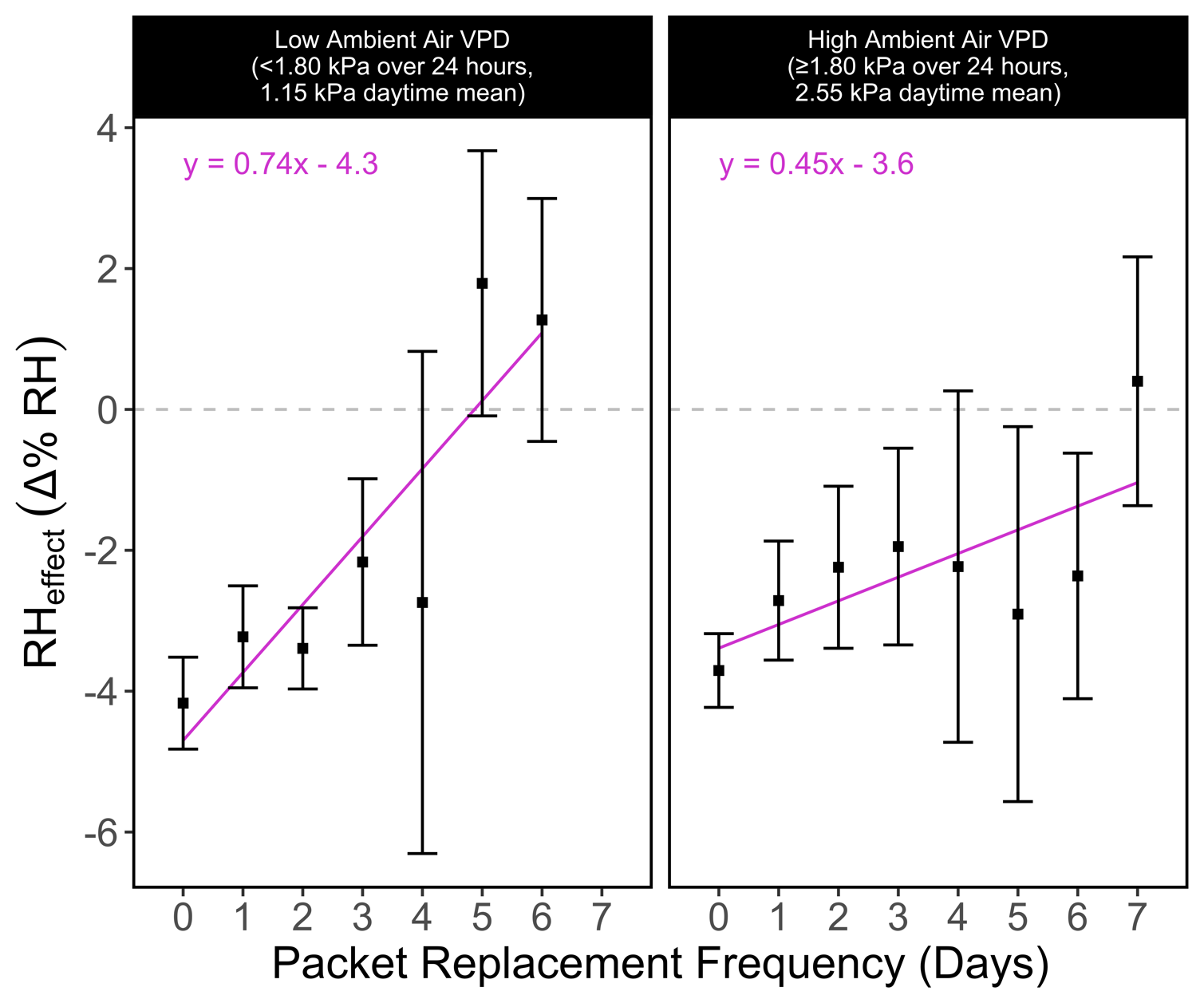

**Figure S2.** We measured changes in RH as a function of packet replacement frequency and average daily VPD. Packets retained longer-term dehumidification capacity (and thus required less frequent replacement) on hot, dry days (of high ambient VPD). Trendlines indicate significant relationships and error bars represent 95% confidence intervals around mean points.

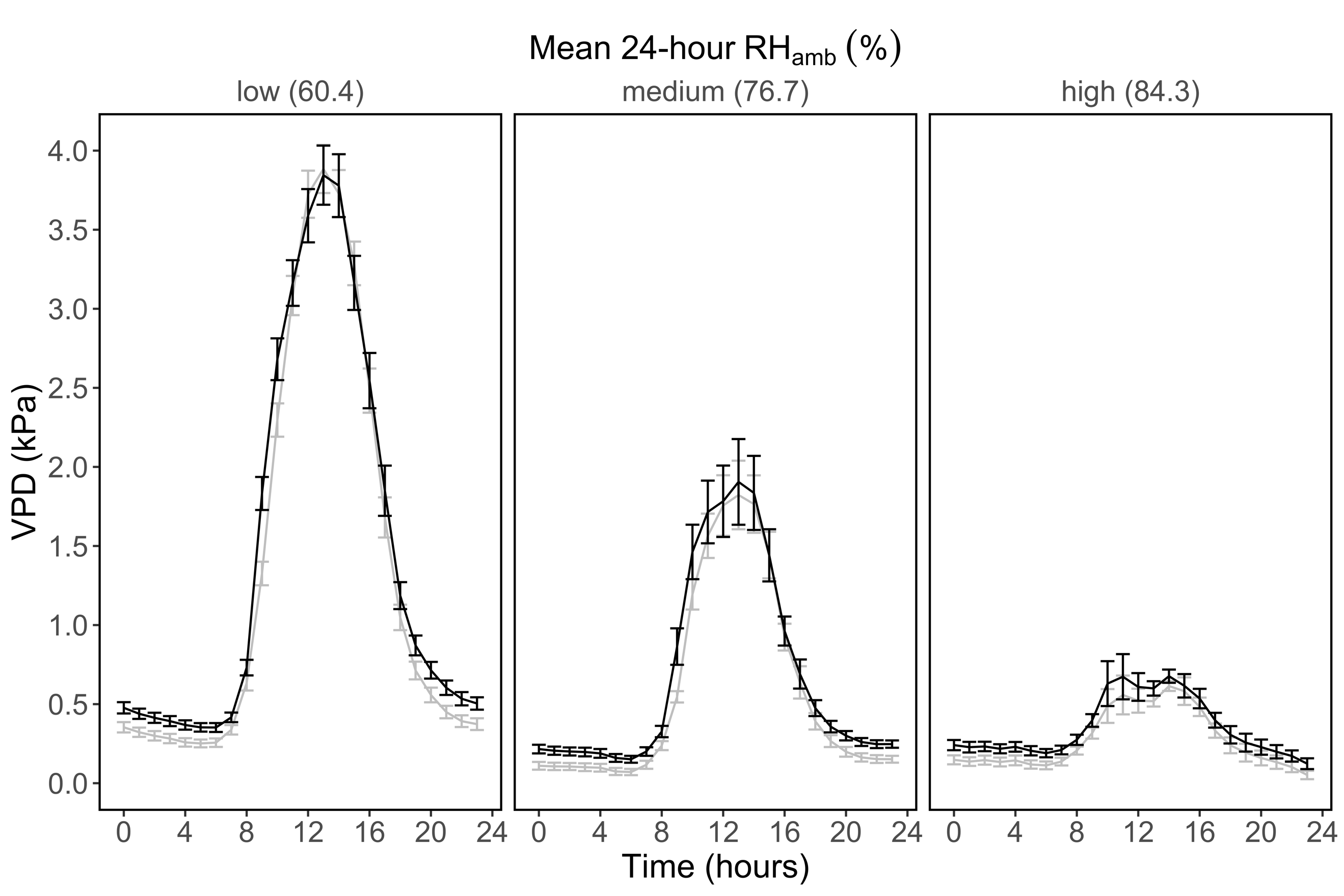

**Figure S3.**  We measured fine-scale changes in VPD over the course of a day. Hourly VPD in pots with silica packets (black lines) can be compared with ambient air VPD in nearby non-treated pots (gray lines). Both varied with respect to ambient air RH (panels). Error bars represent 95% confidence intervals.

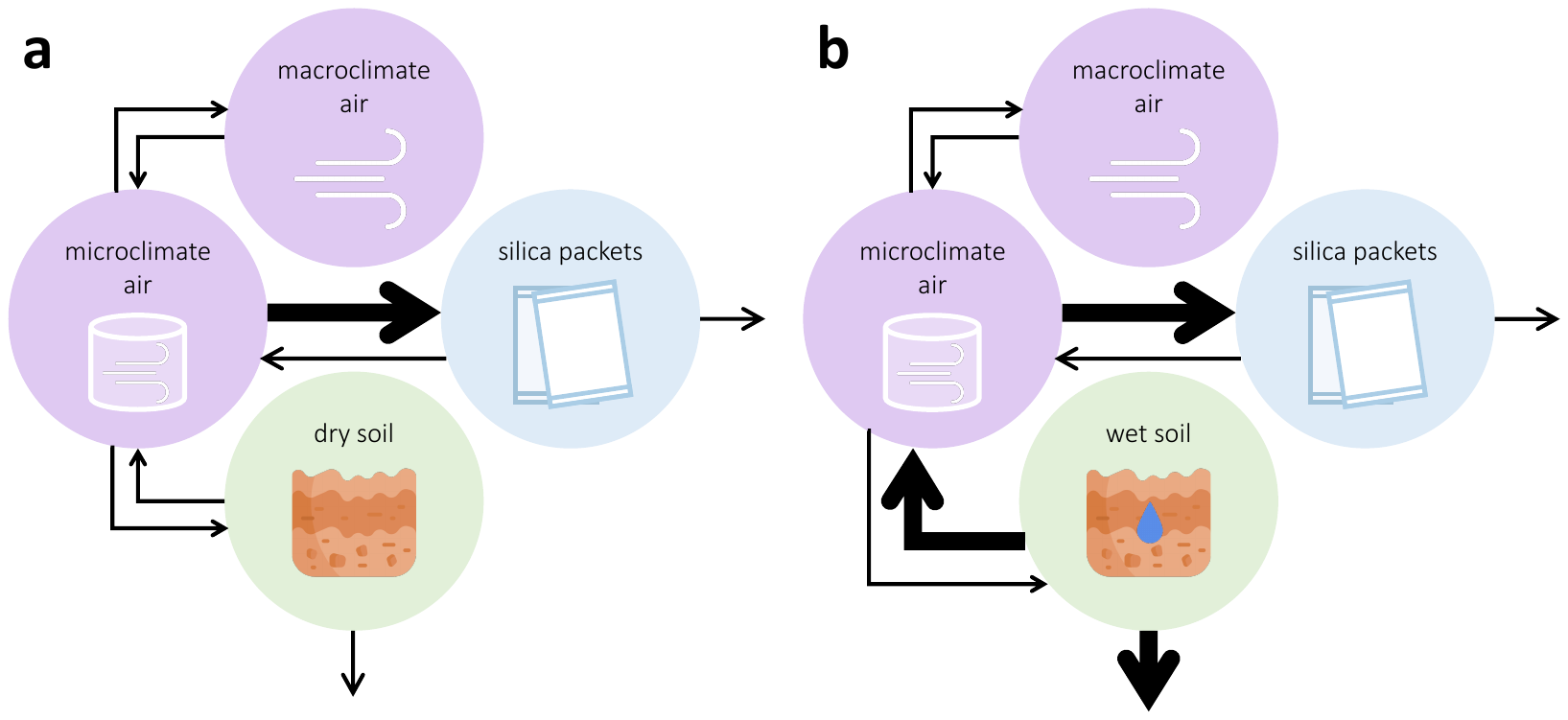
**Figure S4.** In this conceptualization, the movement of water through a miniature water cycle captured by our mesocosms is indicated via arrows between pools. Due to our open-top chamber design, water vapor is exchanged between the macroclimate and microclimate. Water is also absorbed by the soil and evaporates off the surface of the soil. Silica packets can either capture microclimate humidity or re-emit it, depending on water concentration gradients. In this way, soil moisture can drive packet efficiency. In pots of wet soil (**b**), even if packets are capturing humidity, rapid soil moisture evaporation can replace it. In pots of dry soil (**a**), there is less soil moisture evaporation, and thus packets can maintain microclimate humidity at lower levels.

**Protocol for desaturating used packets for redeployment.**

Silica gel packets can be reused repeatedly by drying out the humidity they captured in the field. In our study, after removing saturated packets from our mesocosms, we transferred them to a Fisher Scientific drying oven for approximately 2-3 days. To drive off any remaining moisture, we heated 8-10 packets at a time in a 1250-watt microwave oven (NN-SN966S, Panasonic) on a defrost setting for 15-20 minutes. Ideal dry mass was ≤107.0 g (as determined by warming 10 packets until they stopped losing moisture, and then averaging their individual weights). Once saturated packets were dried to ≤107.0 g, we immediately stored them in airtight plastic bags until redeployment.
